# Points2Regions: Fast, interactive clustering of imaging-based spatial transcriptomics data

**DOI:** 10.1101/2022.12.07.519086

**Authors:** Axel Andersson, Andrea Behanova, Christophe Avenel, Jonas Windhager, Filip Malmberg, Carolina Wählby

## Abstract

Imaging-based spatial transcriptomics techniques generate image data that, once processed, results in a set of spatial points with categorical labels for different mRNA species. A crucial part of analyzing downstream data involves the analysis of these point patterns. Here, biologically interesting patterns can be explored at different spatial scales. Molecular patterns on a cellular level would correspond to cell types, whereas patterns on a millimeter scale would correspond to tissue-level structures. Often, clustering methods are employed to identify and segment regions with distinct point-patterns. Traditional clustering techniques for such data are constrained by reliance on complementary data or extensive machine learning, limiting their applicability to tasks on a particular scale. This paper introduces ‘Points2Regions’, a practical tool for clustering spatial points with categorical labels. Its flexible and computationally efficient clustering approach enables pattern discovery across multiple scales, making it a powerful tool for exploratory analysis. Points2Regions has demonstrated efficient performance in various datasets, adeptly defining biologically relevant regions similar to those found by scale-specific methods. As a Python package integrated into TissUUmaps and a Napari plugin, it offers interactive clustering and visualization, significantly enhancing user experience in data exploration. In essence, Points2Regions presents a user-friendly and simple tool for exploratory analysis of spatial points with categorical labels.

## Introduction

Spatial omics is a rapidly growing field, and several spatial omics techniques are derived from imaging methods, such as multiplexed immunohistochemical staining, cyclic immune fluorescence staining, image-mass cytometry (IMC), and imaging-based spatial transcriptomics (IST) [1]. These omics techniques enable researchers to study the spatial cellular and molecular architecture in tissue, which is crucial for furthering our understanding of biological processes and diseases.

The data obtained from these imaging techniques are primarily in the form of images. However, processing these images makes it common to end up with a point-like output, where each point has a spatial position and categorical label. For example, an IMC experiment would generate high-plex images with protein expression. Analyzing these images makes it possible to identify points indicating the location of different cell types. Likewise, an IST experiment would produce images with fluorescent spots that combinatorically label mRNA molecules. Through decoding, one ends up with points indicating the location of different mRNA types.

Regardless of the type of label, biological regions of interest are usually identified not just by a single point but by a mixture of different types of points in a region. In IST, in particular, groups of mRNA molecules co-localized within a cell can reveal the cell’s function. Likewise, groups mRNAs in a larger ‘niche’ can reveal biological functions on an organ level. A common procedure for identifying these regions is to cluster spatial neighborhoods based on their composition of labels. The size of these neighborhoods depends on the research question but typically ranges from the subcellular scale up to the scale of the tissue composition or even the organ level.

This type of spatial clustering has been widely used within IST, where it is common to cluster spatial neighborhoods corresponding to segmented cells based on their composition of mRNA molecules into clusters corresponding to cell types [2–4, 4–6].

Once the cell types are defined, a second set of niche identification methods comes into play. These methods focus on finding niches by clustering spatial neighborhoods based on their composition of cell types (and/or mRNA molecules) [7–13]. Here, the neighborhoods are large enough to cover multiple cells.

This niche identification type of clustering is relevant to other omics techniques as well. For example, Schurch et al. [14] and Goltsev et al. [15] used an immunofluorescence technique to identify points corresponding to different cell types, and by computing the composition of points in neighborhoods, they identified regions corresponding to cell population niches.

While these cell typing and niche identification methods are powerful, they may face constraints when applied beyond their original context, as they make assumptions about the availability and quality of complementary data, such as cell segmentation [2–9] or single-cell RNA sequencing data [2], thereby restricting their applicability across domains.

An alternative set of methods are cell segmentation-free methods [16–22]. Here, the spatial neighborhoods that are to be clustered are not based on cell segmentation, but instead based on the underlying distribution of points. These methods typically do not rely on complementary data and can potentially be applied in a broader context. A great example is SSAM [17, 18], which was used for identifying structures corresponding to both cell types and niches.

A common trait among all these clustering methods is that they are oriented towards exploratory rather than confirmatory analysis. The aim is to uncover hidden structures and define categories for hypothesis generation rather than quantifying already known patterns. However, for practical exploratory analysis, a crucial criterion is flexibility regarding the ability to capture spatial relations at various scales. The discovery of patterns across various spatial resolutions would benefit from a method that is easy to tune and has interpretable parameters that yield biologically relevant results without lengthy optimization. Moreover, the ability to explore data within its spatial context is essential. Given that biological patterns exist at various scales, static visualization often falls short, emphasizing the necessity of interactive solutions.

In spatial omics analysis, we identified a crucial gap — the lack of a versatile, interactive, and simple tool capable of efficiently clustering points across multiple resolutions without relying on additional data or extensive machine learning. Inspired by cell segmentation-free techniques [17–19], we realized that clustering of label compositions within spatial bins could efficiently reveal biologically relevant regions with very low constraints on both memory and computing.

We present Points2Regions, a tool for clustering spatial categorical biomarkers (points) that work for multiple resolutions and handle varying quantities of points. Points2Regions does not make any assumption regarding the availability of complementary data, making it accessible to a wide range of applications.

Understanding the significance of also visualizing data within its spatial context and capitalizing on the computational efficiency of Points2Regions, we extended its functionality by adding it as an interactive plugin in TissUUmaps [23], a software for visualizing large quantities of points on high-resolution images. Furthermore, we have also made Points2eEgions available as a plugin in Napari, an interactive, multi-dimensional image viewer for Python [24].

Points2Regions is available at https://github.com/wahlby-lab/Points2Regions, as a Napari plugin at https://github.com/wahlby-lab/napari-points2regions, and can be installed from PyPI or conda-forge. Additionally, we offer an online version of Points2Regions at https://tissuumaps.github.io/gallery/#_galleries/0_points2regions.md together with several demos, where users to upload their data, run the clustering, and download the clusters without needing to install anything.

## Material & Methods

### Efficient feature extraction

The idea behind Points2Regions is to aggregate spatial biomarkers with categorical labels (points) in local neighborhoods. The composition of different types of points in a neighborhood, represented as a compositional vector (CV), serves as a feature used for clustering the neighborhoods into regions with similar point compositions. Depending on the application, the user should be able to choose different spatial scales. For example, identifying cell type niches requires aggregating cell type points over hundreds of micrometers, while subcellular mRNA patterns are visible by aggregating molecules on a micrometer scale. To make the extraction of CVs fast and memory efficient, we employ a hierarchical interpolation approach.

First, each point is assigned to a series of spatial bins of varying sizes (representing varying spatial resolutions). We let 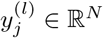 denote the count of the *N* different point-types within the *j*-th bin of the *l*-th resolution level. Each bin’s center position is 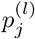. Level *l* = 0 refers to the highest resolution (smallest bin sizes), and *l* = *L* refers to the lowest resolution (largest bin sizes). The grid size, Δ^(*l*)^, increases linearly between the highest and lowest resolution. The CVs are computed at query locations denoted as 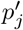 by interpolating the point abundances between the bins using the inverse-distance weighted interpolation:

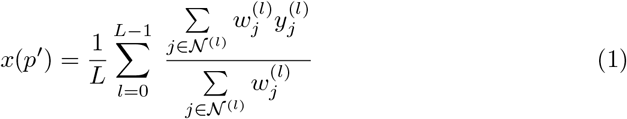

where

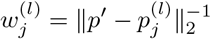

and

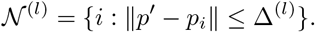

We evaluate Eq. 1 on a rigid grid whose nodes are separated by a distance equal to the width of the bins of the lowest resolution level.

The efficiency of Eq. 1 lies in the swift binning of points onto a grid using the hash function 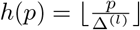 that immediately links a position *p* to a bin index. Moreover, the matrices in Eq. 1 are sparse, allowing for easy storage and multiplication using sparse matrix data structures. Supplementary Fig. 1 shows a 1D example of the CV extraction. We compute the CVs according to Eq 1 before clustering them using mini-batch *k*-means [25].

**Fig 1.**
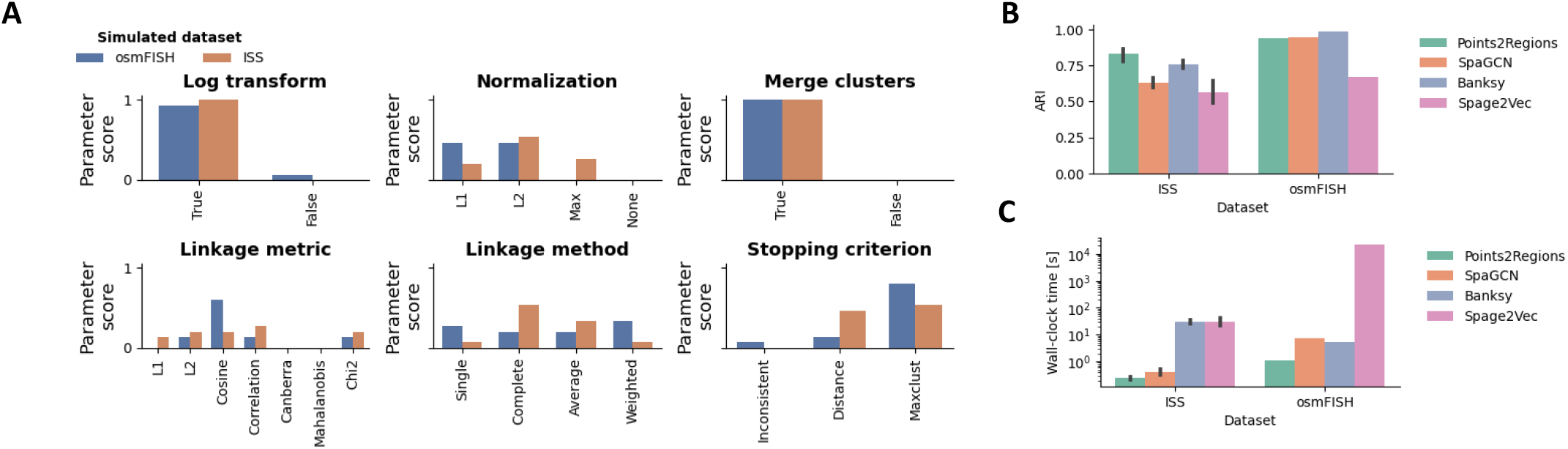
Points2Regions evaluated on simulated data. (**A**) The score for different conditional parameters on simulated data highlights the significance of merging clusters and log transforming CVs. (**B**-**C**) Adjusted rand-index results and corresponding wall-clock times for various methods are presented, highlighting the speed and accuracy of Points2Regions.

### Exploring Synergy in Clustering Methods

While *k*-means clustering is memory-efficient and is generally fast, its limitation is the underlying assumption that clusters are spherical with roughly equal variance. This assumption is not necessarily true. For example, the abundance of a particular biomolecule may vary between different cell types.

Orthogonal to *k*-means clustering is agglomerative (hierarchical) clustering, which can be applied to various data types without assuming its distribution. Hierarchical clustering, however, tends to be slow when the number of observations is large.

Hypothetically, by over-clustering with *k*-means and then merging with hierarchical clustering, we can leverage both the speed of *k*-means and the flexibility of hierarchical clustering, thereby improving the clustering.

There are, however, conditional parameters related to hierarchical clustering, like the *linkage metric* for comparing clusters, the *stopping criterion* that determines when to stop merging the clusters, and the *linkage method* for comparing newly merged clusters. We study a simulated dataset with a known ground truth to empirically determine the best parameters. We also study two additional conditional parameters related to pre-processing: The choice of normalization norm and whether to use log transformation.

### Hyperparameters

Apart from the conditional parameters mentioned above, Points2Regions comes with four tunable hyperparameters:

`Number of clusters`. Determines the number of clusters. It is problem-specific, but several established methods can guide the user [26].

`Bin size`. The width of the lowest resolution bins is also the stride between query locations for computing CVs. It is a trade-off between spatial resolution and computational speed. A larger bin size yields faster clustering but may result in pixelated regions.

`Bin smoothing`. Governs the spatial distance over which points are aggregated to a query location. The width of the lowest resolution bins is set to this value times the pixel size. The value of this parameter is problem-specific. However, it needs to be large enough to ensure a sufficient number of points are included in the CVs so that a co-varying structure can be observed among them. This can be determined by (i) computing the CVs by incrementally varying the pixel smoothing sizes, (ii) conducting principal component analysis with a fixed number of components, and (iii) tracking the explained variance as a function of smoothing distance. The knee-point on this curve indicates when co-varying structures start to appear, and a sufficiently large smoothing distance is used.

`Minimum number of markers per bin`. This is a simple density threshold parameter used for discarding query locations for CVs where the number of markers per lowest-resolution pixels is less than this value.

#### Experiments

### Simulated data for model evaluation

To quantify the effect of different conditional hyperparameters, we simulate two datasets.

First, we consider the simulated osmFISH dataset used in the clustering benchmarking study by Cheng et al. [27]. Here, ground-truth labels corresponding to layers in a mouse brain somatosensory cortex were assigned to individual cells whose location and mRNA content are based on information from the real data [28].

Since segmentation-free methods operate on points for individual mRNA species instead of count-per-cell vectors, we randomly place each mRNA point within radii from the cell center. Cell radii varied randomly between 5 [um] and 20 [um] between different cells.

Next, we use pciSeq, a probabilistic method for assessing IST data, to simulate an *in situ* sequencing dataset. The pciSeq method attempts to assign points for different mRNA species to detected cells and define the cell type by utilizing complementary single-cell RNA sequencing data. The assignment of molecules to cells and the matching of cells to cell types are quantified as probabilities. By setting the probability threshold for an mRNA belonging to a cell to 0.85 and the probability threshold for a cell matching a cell type to 0.85, we identify cleanly segmented and typed cells in the data. These cells should have mRNA compositions that align well with single-cell RNA sequencing data. We use these cells to identify the expected molecular composition for each cell type. As a final step, we go through all cells segmented and typed by Qian et al [2]. If a cell was typed with a probability greater than 0.85, we simulate a new distribution of mRNA molecules using the expected molecular composition of that cell. This process enables us to create a dataset with clean cell types, free from outliers resulting from inadequate segmentation. This dataset was generated from a mouse brain coronal section (hippocampal CA1 region). There were 14 sections in total. For each section, we used both the left and right hippocampal CA1 region, resulting in 28 replicated ISS datasets.

#### Selecting optimal conditional parameters

The simulated datasets offer a well-defined ground truth, which allows us to explore the impact of different conditional parameters used in the method. These parameters are (i) the type of normalization norm of CV, (ii) whether to use log-transformation or not, (iii) whether to use the hierarchical merging or not, (iv) which *linkage metric* to use for comparing clusters, (v) the *linkage method* for calculating the distance between the newly merged clusters, (vi) and the *stopping criterion* that determines when to stop the hierarchical merging of clusters. These conditional parameters are further detailed in Supplementary Text. 2.

To assess performance, we use the adjusted rand-index (ARI) and Optuna [31], a stochastic hyper-parameter tuning package. We run 50 Optuna parameter searches (each with 100 iterations and different seeds) and search for the parameter configuration that results in the highest ARI on the simulated osmFISH and ISS dataset. For the ISS dataset, we chose the replica with the most points. The relative frequency of Optuna searches ending on a particular parameter indicates that parameter’s score. We choose the best-scoring parameters in the final implementation.

### Comparing against prior art

After deciding on the best parameter configuration, we compare Points2Regions against established methods. In the simulated osmFISH dataset, cells are labeled by niches, while in the ISS dataset, individual cells are labeled by type. Therefore, we compare Points2Regions with both niche identification methods [4, 8] and cell typing methods [4, 21]. We optimize hyperparameters using Optuna [31] for all methods, except for [21], where parameters were manually tuned due to lengthy optimization. Additional details are provided in Supplementary Text 1.

#### Experiments on real data

We extend our analysis to four real datasets: a MERFISH dataset by [29], osmFISH dataset [28], Xenium dataset [30], and a MERFISH subcellular dataset [11], and compare Points2Regions against published work. Statistical information for these datasets is available in Table 1.

**Table 1.** Dataset statistics.

| Dataset | Target | # of cells | Avg. # of points per cell | # of mRNA species | Reference |
| --- | --- | --- | --- | --- | --- |
| MERFISH | Niches | 7000 | 250 | 135 | Moffitt et al. [29] |
| osmFISH | Niches & cell types | 5800 | 110 | 35 | Codeluppi et al [28] |
| Xenium | Cell types | 13800 | 270 | 248 | Janesick et al. [30] |
| Merfish sub. | Subcellular structures | 15 | 221090 | 135 | Mah et al. [11] |

## Results

### Parameter optimization highlights the importance of normalization, log-transformation, and cluster merging

First, we ran several parameter searches with Optuna to choose the best conditional parameters. Fig 1A shows the parameter score, that is, the relative frequencies of parameter searches selecting a particular parameter. We noted that almost all models converged to a parameter configuration using log transform, normalization, and hierarchical cluster merging. However, we did not notice a significant difference between which normalization norm to use. We also noted that most of the parameter searches landed in a parameter configuration, including the merging of clusters as post-processing. Based on this analysis, we deterministically set Points2Regions’ linkage metric to cosine distance, its linkage method to complete, and the stopping criterion to `maxclust`. As described in Supplementary Text 2, this stopping criteria means merging clusters until a given number of maximum clusters are obtained. The maximum number of clusters is decided by the user. The number of *k*-means clusters is set to 3*/*2 times this value.

### Benchmarking Points2Regions on simulated data

Next, we compare Points2Region to previously established methods. We consider SpaGCN [8], Banksy [4], Spage2Vec [21]. Fig 1B-C shows the ARI and wall-clock time for the respective methods on the simulated datasets. Points2Region performs similarly to the established methods w.r.t ARI on the osmFISH dataset and remarkably well on the ISS dataset, even though it does not use the cell borders generated when simulating it.

### Points2Regions on real data

We tested Points2Regions on four real image-based datasets for spatial transcriptomics to see if it finds clusters like the original studies. Hyperparameters are presented in Supplementary Text 1. All of the Points2Regions clusters can be interactively explored at https://tissuumaps.dckube.scilifelab.se/web/private/Axel/points2regions.html.

We first study the MERFISH dataset [32]. We chose this dataset since it is used to evaluate the popular niche-identification method SpaGCN [8]. Here, we want to see if Points2Regions can, without relying on cell segmentation, identify a region similar to SpaGCN. Fig. 2A shows a visual comparison between the clusters identified by Points2Regions and SpaGCN. SpaGCN is used on the mRNA transcripts within segmented cells, whereas Points2Regions runs all mRNA points. Additionally, Fig. 2B presents the relative frequency of each cluster identified by Points2Regions across the different SpaGCN clusters. A notable observation from the study is that Points2Regions successfully captured many of the larger structures in the dataset. Specifically, eight out of the ten identified clusters showed a strong correspondence with those identified by SpaGCN, as illustrated in Fig. 2B. The missing two clusters were of the lowest abundance and likely do not correspond to the niche of cell types. This highlights the effectiveness of Points2Regions in uncovering biologically significant information without the necessity for cell segmentation.

**Fig 2.**
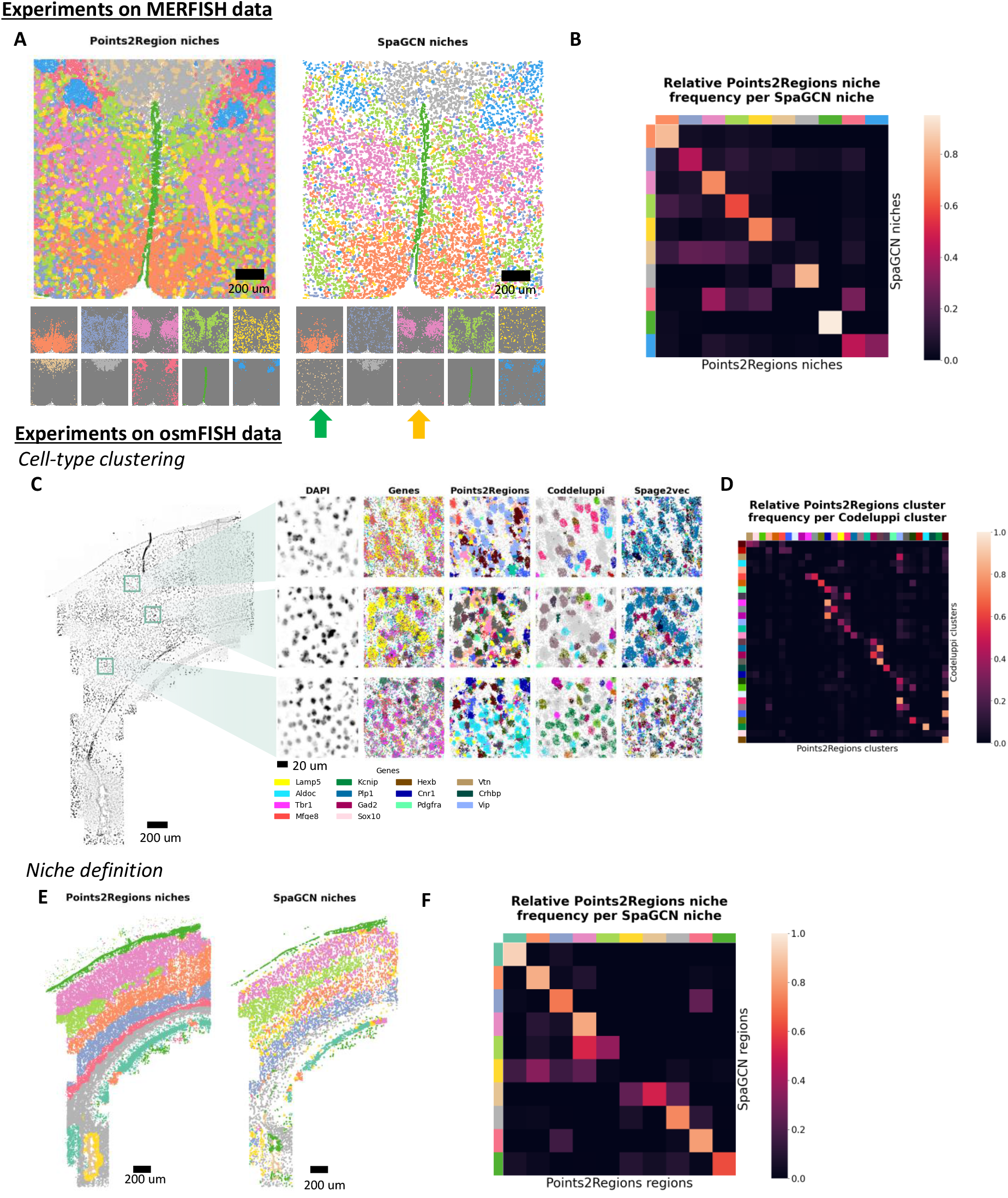
Points2Regions breaks down two large spatial transcriptomics datasets. (**A**) Comparison of clusters defined by Points2Regions and SpaGCN on the MERFISH dataset. Segmented cells are grouped into niches using SpaGCN, and regions are defined by Points2Region without any cell-segmentation. The green and orange arrows show the niches identified by SpaGCN that did not correspond well with any Points2Reions niches. (**B**) The distribution of Points2Region clusters for each SpaGCN cluster. Despite Points2Regions operating directly on transcripts without cell segmentation, both methods reveal similar clusters. (**C**) Points2Regions clusters on osmFISH data are shown with original clusters by Codeluppi et al. [28] and Spage2Vec [21]. Gray points represent excluded points in the clustering. (**D**) Distribution of Points2Regions clusters for each Codeluppi cluster. (**F**-**G**) Comparison of identified niches using Points2Regions (large smoothing distance) and SpaGCN.

Next, we studied the osmFISH dataset. Fig 2C shows clusters from Points2Regions, the original study (Codeluppi et al. [28]) and Spage2vec, on the osmFISH dataset. Codeluppi clusters were found by segmenting cells and then clustering, while Points2Regions and Spage2vec work directly on the transcripts. Comparing the gene distribution in Fig 2C with the Points2Regions clusters, we note that Points2Regions tends to pick up on clusters corresponding to cell types. Fig 2D compares the distribution of Points2Regions clusters per Codeluppi cluster. A perfect diagonal would indicate a one-to-one mapping between the methods. Fig 2D also indicates a correlation between Points2Regions clusters and cell clusters defined by Codeluppi et al [28].

We also ran SpaGCN on the osmFISH dataset to identify niches corresponding to different cortical layers in the brain and compare those with niches defined using Points2Region. Here, we ran Points2Regions with a much larger bin smoothing, and we can observe a correlation between SpaGCN and Points2Regions, see Fig 2E-F. Despite being simple and efficient, Points2Regions can identify structures similar to established scale-speficic pipelines.

Next, we studied the Xenium dataset and compared Points2Regions clusters with those found by Xenium’s graph-based clustering. We also ran Points2Regions with increasing bin sizes and smoothing distances to showcase its utility in identifying patterns across different scales. The clustering of the highest resolution took roughly two minutes to complete on a regular laptop, showcasing its efficiency on a dataset with over 40M points. The resulting clusters exhibited a notable similarity to those provided by the Xenium instrument, see Fig 3A-C, reinforcing Points2Regions’ consistency and reliability. Supplementary Fig. 2 compares the Points2Regions niche-level clusters (right-most figure) with a reference image from Allen Brain Atlas [33, 34]. We noticed that Points2Regions identified several known compartments in the mouse brain. Most evidently are the layers in the somatosensory area.

**Fig 3.**
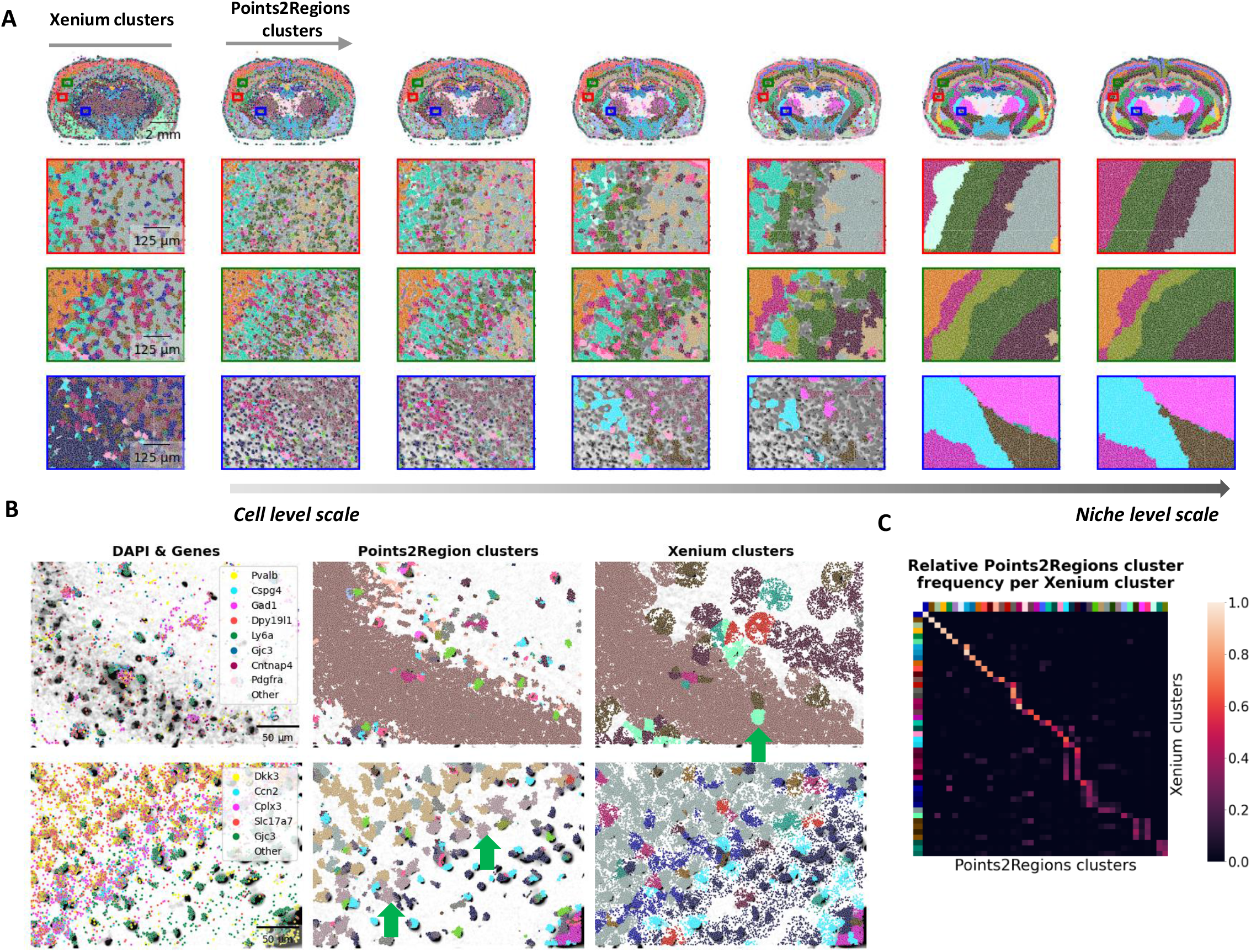
Application of Points2Region on data generated by state-of-the art *in situ* sequencing instrument (Xenium). (**A**) Comparison between Points2Region and clusters produced by the Xenium software. The leftmost column shows Xenium clusters. The other columns show Points2Region clusters obtained by aggregating points over incrementally larger distances. (**B**) Two zoomed-in regions with genes, Points2Region clusters and alongside Xenium clusters. Green arrows mark clusters that, based on the gene expression, are successfully identified by one method and not by the other. (**C**) The relative frequency of Points2Region clusters for each Xenium cluster.

Finally, we considered a MERFISH dataset with a remarkably large number of mRNA points per cell. We ran Points2Regions with four clusters to see if we could identify sub-cellular structures similar to Mah et al. [11]; we noticed several similarities. For example, nuclei-septic clusters have high abundances of *MALAT1*, cytoplastic-specific clusters have high abundances of *TLN1*, and nuclear-edge-specific clusters have high abundances of *FBN2*, see Fig. 4. This observation highlights Points2Regions’ versatility, showcasing its applicability to various tasks.

**Fig 4.**
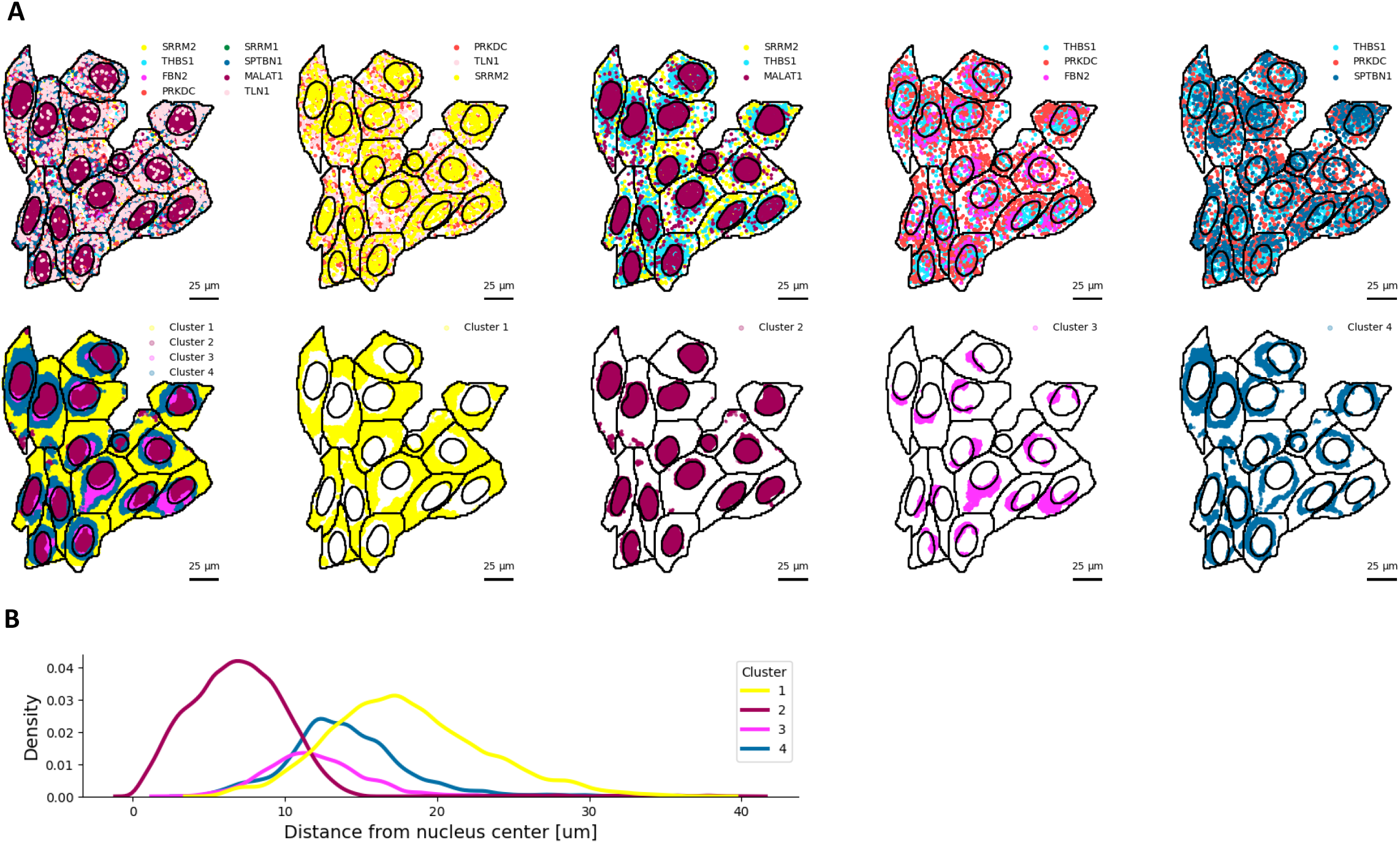
Application of Points2Regions on high-resolution image-based spatial transcriptomics data, revealing subcellular structures. (**A**) Highly abundant genes along Points2Regions results (bottom) alongside manual annotations (black lines) from the original study [27]. (**B**) Illustration of cluster density as a function of distance to the nuclei center.

## Discussion

We introduced Points2Regions, a fast and interactive tool for clustering and delineating regions in categorical marker data commonly encountered in spatial omics. Unlike many methods relying on predefined cells or complementary single-cell data, our approach embraces a straightforward, segmentation-free method. Rather than aiming for the most powerful methods, we aim to create an algorithm that is effective, fast, and easy-to-execute across different scales. Points2Regions performs remarkably well on simulated datasets. On the osmFISH simulated dataset, both Banksy and SpaGCN perform only slightly better than Points2Regions, even though they rely on perfectly segmented cells, while Points2Regions does not use any information from cell segmentation. Moreover, Points2Regions excels on the ISS dataset, outperforming other methods. Here, the objective is individual cell typing rather than identifying cell type niches. Therefore, the global spatial context that SpaGCN and Banksy facilitate is less relevant, which could explain their poor performance. It should be noted, though, that SpaGCN is designed for niche definition rather than cell-typing. Moreover, the number of cells and reads per cell is lower for the simulated ISS dataset, which makes it harder for unsupervised deep learning employed by SpaGCN.

We observed notable similarities when comparing clusters generated by Points2Regions with those produced by other methods using real data. This suggests that Points2Regions is adept at identifying biologically relevant features efficiently. We observed instances where Points2Regions failed to detect a region identified by another method or vice versa. See the green arrow in Fig. 3 or green and orange arrow in Fig. 2. In such cases, a relatively small adjustment of hyperparameters often facilitated the discovery of the missing cell type. However, establishing a one-to-one mapping between clustering methods proves challenging, considering the diverse algorithmic underlying assumptions made by each method.

In our experiments, merging *k*-means clusters improved clustering performance. All parameter searches on simulated data ended up with cluster merging turned on. The hierarchical clustering might mitigate some underlying assumptions of *k*-means, such as the assumption that spherical clusters have similar variance. We explored only the hierarchical options available in SciPy [35, 36], but other methods like those in Peterson et al. [37] could be interesting.

However, a drawback of the “counting points over spatial distances” approach is its insensitivity to edges. Sharp boundaries in spatial biology often separate different regions with distinct point compositions. Counting points over too large a distance may mix points from two regions, leading to potential misinterpretation. Exploring edge-preserving aggregation functions inspired by filters in image analysis, such as bilateral filtering [38] or image restoration via graph cuts [39], could address this issue.

To conclude, Points2Regions provides an efficient interpretation of large and diverse spatial omics datasets. When combined with interactive visualization via TissUUmaps or Napari, Points2Regions can increase our understanding of biological systems at various spatial resolutions.

## Data Availability

The analyzed datasets were acquired from the following websites:

- The simulated osmFISH dataset [27] is available at https://github.com/acheng416/Benchmark-CTCM-ST.
- The simulated ISS dataset is available at `URL`.
- The osmFISH dataset is available by the authors [28] at https://linnarssonlab.org/osmFISH/availability/.
- The MERFISH dataset is available by the authors [29] at https://datadryad.org/stash/dataset/doi:10.5061/dryad.8t8s248.
- Xenium dataset is provided by 10x Genomics and is available at https://www.10xgenomics.com/resources/datasets/fresh-frozen-mouse-brain-for-xenium-explorer-demo-1-standard.

## Code availability

- Points2Regions can be installed as a Python package from https://github.com/whalby-lab/Points2Regions.
- Points2Regions can be installed as a TissUUmaps plugin inside the TissUUmaps software [23].
- Points2Regions can be installed as a Napari plugin from https://github.com/wahlby-lab/napari-points2regions.
- Points2Regions is available as on online ‘uplooad-data-and-download-regions’ service here: https://tissuumaps.github.io/gallery/#_galleries/0_points2regions.md.

## Notes

### Competing Interest Statement

The authors have declared no competing interest.

### Summary of Updates

The manuscript has been reworked

## References

1. Bressan D, Battistoni G, Hannon GJ. The dawn of spatial omics. Science. 2023;381(6657):eabq4964.

2. Qian X, Harris KD, Hauling T, Nicoloutsopoulos D, Muñoz-Manchado AB, Skene N, et al. Probabilistic cell typing enables fine mapping of closely related cell types in situ. Nature methods. 2020;17(1):101–106.

3. Prabhakaran S. Sparcle: assigning transcripts to cells in multiplexed images. Bioinformatics Advances. 2022;2(1):vbac048.

4. Singhal V, Chou N, Lee J, Liu J, Chock WK, Lin L, et al. BANKSY: a spatial omics algorithm that unifies cell type clustering and tissue domain segmentation. bioRxiv. 2022; p. 2022–04.

5. Wolf FA, Angerer P, Theis FJ. SCANPY: large-scale single-cell gene expression data analysis. Genome biology. 2018;19:1–5.

6. Chen JG, Chávez-Fuentes JC, O’Brien M, Xu J, Ruiz E, Wang W, et al. Giotto Suite: a multi-scale and technology-agnostic spatial multi-omics analysis ecosystem. bioRxiv. 2023; p. 2023–11.

7. Yu N, Zhang D, Zhang W, Liu Z, Qiao X, Wang C, et al. stGCL: A versatile cross-modality fusion method based on multi-modal graph contrastive learning for spatial transcriptomics. bioRxiv. 2023; p. 2023–12.

8. Hu J, Li X, Coleman K, Schroeder A, Ma N, Irwin DJ, et al. SpaGCN: Integrating gene expression, spatial location and histology to identify spatial domains and spatially variable genes by graph convolutional network. Nature methods. 2021;18(11):1342–1351.

9. Teng H, Yuan Y, Bar-Joseph Z. Clustering spatial transcriptomics data. Bioinformatics. 2022;38(4):997–1004.

10. El Marrahi A, Lipreri F, Kang Z, Gsell L, Eroglu A, Alber D, et al. NIPMAP: niche-phenotype mapping of multiplex histology data by community ecology. Nature Communications. 2023;14(1):7182.

11. Mah CK, Ahmed N, Lopez N, Lam D, Monell A, Kern C, et al. Bento: A toolkit for subcellular analysis of spatial transcriptomics data. BioRxiv. 2022; p. 2022–06.

12. Yuan Z. MENDER: fast and scalable tissue structure identification in spatial omics data. Nature Communications. 2024;15(1):207.

13. Hu Y, Rong J, Xu Y, Xie R, Peng J, Gao L, et al. Unsupervised and supervised discovery of tissue cellular neighborhoods from cell phenotypes. Nature Methods. 2024; p. 1–12.

14. Schürch CM, Bhate SS, Barlow GL, Phillips DJ, Noti L, Zlobec I, et al. Coordinated cellular neighborhoods orchestrate antitumoral immunity at the colorectal cancer invasive front. Cell. 2020;182(5):1341–1359.

15. Goltsev Y, Samusik N, Kennedy-Darling J, Bhate S, Hale M, Vazquez G, et al. Deep profiling of mouse splenic architecture with CODEX multiplexed imaging. Cell. 2018;174(4):968–981.

16. Petukhov V, Xu RJ, Soldatov RA, Cadinu P, Khodosevich K, Moffitt JR, et al. Cell segmentation in imaging-based spatial transcriptomics. Nature biotechnology. 2022;40(3):345–354.

17. Park J, Choi W, Tiesmeyer S, Long B, Borm LE, Garren E, et al. Cell segmentation-free inference of cell types from in situ transcriptomics data. Nature communications. 2021;12(1):3545.

18. Tiesmeyer S, Sahay S, Müller-Bötticher N, Eils R, Mackowiak SD, Ishaque N. SSAM-lite: a light-weight web app for rapid analysis of spatially resolved transcriptomics data. Frontiers in Genetics. 2022;13:785877.

19. Partel G, Hilscher MM, Milli G, Solorzano L, Klemm AH, Nilsson M, et al. Automated identification of the mouse brain’s spatial compartments from in situ sequencing data. BMC biology. 2020;18(1):1–14.

20. Si Y, Lee C, Hwang Y, Yun JH, Cheng W, Cho CS, et al. FICTURE: Scalable segmentation-free analysis of submicron resolution spatial transcriptomics. bioRxiv. 2023; p. 2023–11.

21. Partel G, Wählby C. Spage2vec: Unsupervised representation of localized spatial gene expression signatures. The FEBS Journal. 2021;288(6):1859–1870.

22. Walter FC, Stegle O, Velten B. FISHFactor: a probabilistic factor model for spatial transcriptomics data with subcellular resolution. Bioinformatics. 2023;39(5):btad183.

23. Pielawski N, Andersson A, Avenel C, Behanova A, Chelebian E, Klemm A, et al. TissUUmaps 3: Improvements in interactive visualization, exploration, and quality assessment of large-scale spatial omics data. Heliyon. 2023;9(5).

24. Ahlers J, Althviz Moré D, Amsalem O, Anderson A, Bokota G, Boone P, et al. napari: a multi-dimensional image viewer for Python; 2023. Available from: https://zenodo.org/record/3555620.

25. Pedregosa F, Varoquaux G, Gramfort A, Michel V, Thirion B, Grisel O, et al. Scikit-learn: Machine Learning in Python. Journal of Machine Learning Research. 2011;12:2825–2830.

26. Schubert E. Stop using the elbow criterion for k-means and how to choose the number of clusters instead. ACM SIGKDD Explorations Newsletter. 2023;25(1):36–42.

27. Cheng A, Hu G, Li WV. Benchmarking cell-type clustering methods for spatially resolved transcriptomics data. Briefings in Bioinformatics. 2023;24(1):bbac475.

28. Codeluppi S, Borm LE, Zeisel A, La Manno G, van Lunteren JA, Svensson CI, et al. Spatial organization of the somatosensory cortex revealed by osmFISH. Nature methods. 2018;15(11):932–935.

29. Moffitt JR, Bambah-Mukku D, Eichhorn SW, Vaughn E, Shekhar K, Perez JD, et al. Molecular, spatial, and functional single-cell profiling of the hypothalamic preoptic region. Science. 2018;362(6416):eaau5324.

30. Janesick A, Shelansky R, Gottscho A, Wagner F, Rouault M, Beliakoff G, et al. High resolution mapping of the breast cancer tumor microenvironment using integrated single cell, spatial and in situ analysis of FFPE tissue. BioRxiv. 2022; p. 2022–10.

31. Akiba T, Sano S, Yanase T, Ohta T, Koyama M. Optuna: A next-generation hyperparameter optimization framework. In: Proceedings of the 25th ACM SIGKDD international conference on knowledge discovery & data mining; 2019. p. 2623–2631.

32. Moffitt JR, Bambah-Mukku D, Eichhorn SW, Vaughn E, Shekhar K, Perez JD, et al. Molecular, spatial, and functional single-cell profiling of the hypothalamic preoptic region. Science. 2018;362(6416):eaau5324.

33. Allen Institute for Brain Science. Allen Mouse Brain Atlas; 2004. http://mouse.brain-map.org.

34. Allen Institute for Brain Science. Allen Reference Atlas – Mouse Brain; 2011. http://atlas.brain-map.org.

35. SciPy v1.11.4 Manual;. Available from: https://docs.scipy.org/doc/scipy/reference/generated/scipy.cluster.hierarchy.fcluster.html#scipy.cluster.hierarchy.fcluster.

36. Virtanen P, Gommers R, Oliphant TE, Haberland M, Reddy T, Cournapeau D, et al. SciPy 1.0: fundamental algorithms for scientific computing in Python. Nature methods. 2020;17(3):261–272.

37. Peterson AD, Ghosh AP, Maitra R. Merging K-means with hierarchical clustering for identifying general-shaped groups. Stat. 2018;7(1):e172.

38. Tomasi C, Manduchi R. Bilateral filtering for gray and color images. In: Sixth international conference on computer vision (IEEE Cat. No. 98CH36271). IEEE; 1998. p. 839–846.

39. Boykov Y, Veksler O, Zabih R. Fast approximate energy minimization via graph cuts. IEEE Transactions on pattern analysis and machine intelligence. 2001;23(11):1222–1239.

