## Supplementary Figures for "Points2Regions: Fast, interactive clustering of imaging-based spatial transcriptomics data"

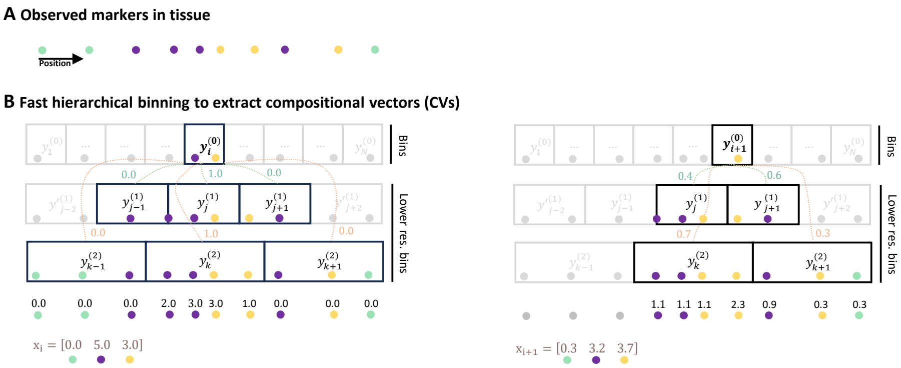


**SF Fig. 1.** Graphical 1D explanation of the hierarchical binning. Categorical markers are observed in the tissue (**A**) and counted in bins of increasing sizes (**B**). The compositional vector, $x_i$, computed at a particular query location, is obtained by weighing together all bins that are a distance less than the bin width from the query location. The weighing is proportional to the inverse distance between the query location and the centre of the bins.

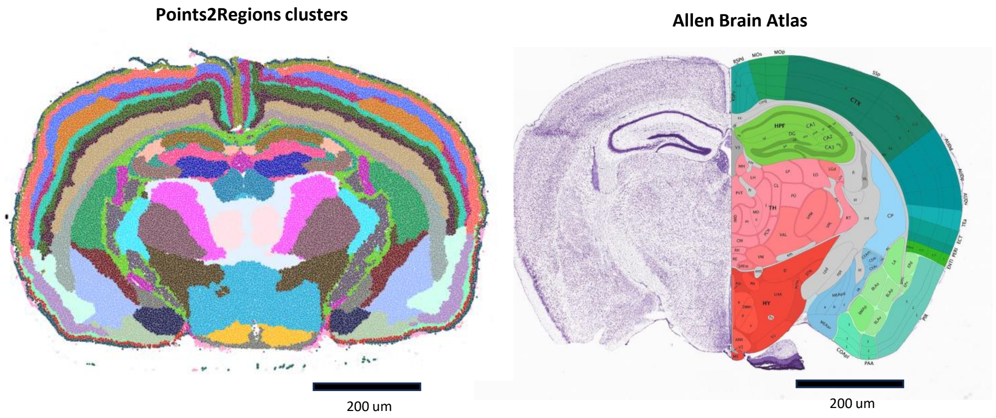


**SF Fig. 2.** Comparison between Points2Regions clusters from the Xenium mouse brain dataset and and anatomical annotations and Nissl staining from the Allen Mouse Brain Atlas and Allen Reference Atlas. The Points2Regions clusters reveal many compartment-level structures in the brain, such as the layers in the somatosensory area.
