## Supplementary Text for "Points2Regions: Fast, interactive clustering of imaging-based spatial transcriptomics data"

### 1 Hyperparameter tunig

#### Optimal paramters for the simulated datasets

We employed Optuna [1], a hyperparameter tuning framework, to search the optimal parameters for the different models. Due to the lengthy optimization time, Spage2Vec was manually tuned on the simulated osmFISH dataset. For the remaining methods, we ran Optuna for 500 iterations. Given the varying number of target clusters across different ISS replicas, each model was separately tuned for each replica. The best-tuned parameters for each replica were then used for assessing the model performance using the adjusted rand index.

**ST Table. 1.** The table contains hyperparameter tuned using Optuna [1] for the benchmarking of different methods on the two simulated datasets. The parameters were tuned by running Optuna for 500 iterations. The parameters were tuned manually for Spage2Vec, due to lengthy optimization time.

| Method | Parameter | Data type | Range |
|---|---|---|---|
| `Points2Regions` | # of clusters | Integer | [3, 50] |
| | Pixel size | Float | [0.1, 20] |
| | Smoothing | Float | [1, 30] |
| | Density threshold | Float | [0, 50] |
| `SpaGCN` | Min cells | Integer | [0, 20] |
| | p | Float | [0, 1.0] |
| | Resolution | Float | [0.1, 2.0] |
| | NN | Integer | [1, 100] |
| | Num PCs | Integer | [1, 75] |
| `Banksy` | Max labels | Integer | [3, 50] |
| | Resolution | Float | [0.1, 2.0] |
| | PCA dimensions | Integer | [1, 75] |
| | Lam | Float | [0, 1] |
| | k geom | Integer | [1, 30] |
| | Weight decay | Categorical | ['scaled_gaussian', 'reciprocal'] |
| | Add non-spatial | Categorical | [True, False] |
| | Variance balance | Categorical | [True, False] |
| | Min cells | Integer | [0, 20] |
| | Normalize per cell | Categorical | [True, False] |
| | Log transform | Categorical | [True, False] |
| `Spage2Vec (osmFISH)` | % | Float | 99 |
| | Num neg samples | Integer | 30 |
| | Walk length | Integer | 2 |
| | NN | Integer | 14 |
| | Resolution | Float | 0.15 |

### Points2Region parameters on the real dataset

Table 2 contains Points2Region parameters for the different datasets considered in the study.

**ST Table. 2.** Points2Region parameters on the four real datasets used in the study.

| **Dataset** | **# of clusters** | **Pixel size** [um] | **Smoothing** [pixels] | **Density threshold** | **Reference** |
|---|---|---|---|---|---|
| osmFISH (cell types) | 31 | 1 | 5 | 5 | [2] |
| osmFISH (niches) | 10 | 1 | 50 | 5 | [2] |
| MERFISH | 10 | 1 | 20 | 40 | [3] |
| Xenium (level 1) | 42 | 1 | 5 | 10 | [4] |
| Xenium (level 2) | 42 | 2 | 5 | 40 | [4] |
| Xenium (level 3) | 42 | 4 | 5 | 160 | [4] |
| Xenium (level 4) | 42 | 8 | 5 | 320 | [4] |
| Xenium (level 5) | 42 | 8 | 10 | 320 | [4] |
| Xenium (level 6) | 42 | 8 | 15 | 320 | [4] |
| MERFISH sub cell. | 4 | 0.1 | 35 | 5 | [5] |

### 2 Conditional parameters

The different conditional parameters are listed in Fig. 1. The conditional parameters are:

`Log-transform`: Whether or not the compostional vectors (CVs) should be transformed using the $\log(x+1)$ function.

`Normalization`: Which norm to use when normalizing the computed CVs.

`Linkage metric`: Which metric to use for comparing the clusters in the hierarchical merging of clusters. We consider the metrics: L2, L1, Cosine, Correlation, Canberra, Mahalanobis and Chi2.

`Linkage formula`: Which formula to consider when comparing newly merged clusters. See [6] for mathematical description.

`Stopping criterion`: Which criterion to use for determining when to stop the cluster merging. We investigate three stopping critera [7, 8]:

- `Inconsistent`: If a node in the dendrogram and all its children have an inconsistent value less than or equal to $t$, then all children belong to the same merged cluster. This is repeated until no more clusters can be merged.
- `Distance`: Merge the $k$-means clusters so that the original $k$-means clusters in each merged cluster have a cophenetic distance less than $t$.
- `Maxclust`: Finds a minimum threshold $r$ so that the cophenetic distance between any two original $k$-means clusters in the same merged cluster is no more than $r$ and no more than $t$ merged clusters are formed.

See SciPy documentation for details [7, 8].

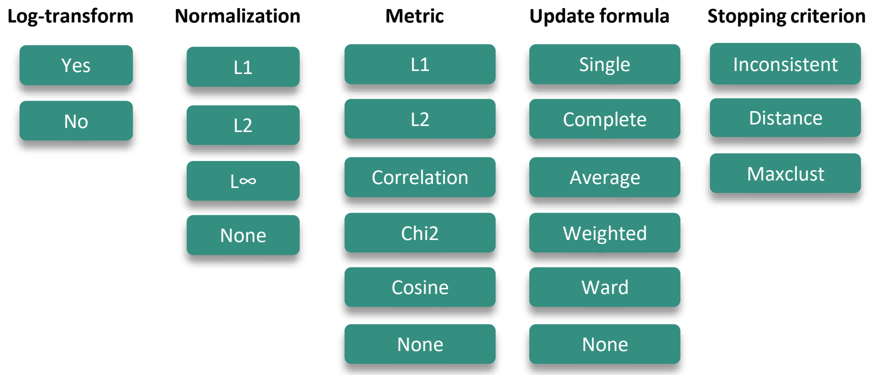


**ST Fig. 1.** Different conditional parameter choices investigate in the parameter study on the simulated data.

### References


1. Akiba T, Sano S, Yanase T, Ohta T, Koyama M. Optuna: A next-generation hyperparameter optimization framework. In: Proceedings of the 25th ACM SIGKDD international conference on knowledge discovery & data mining; 2019. p. 2623–2631.

2. Codeluppi S, Borm LE, Zeisel A, La Manno G, van Lunteren JA, Svensson CI, et al. Spatial organization of the somatosensory cortex revealed by osmFISH. Nature methods. 2018;15(11):932–935.

3. Moffitt JR, Bambah-Mukku D, Eichhorn SW, Vaughn E, Shekhar K, Perez JD, et al. Molecular, spatial, and functional single-cell profiling of the hypothalamic preoptic region. Science. 2018;362(6416):eaau5324.

4. Janesick A, Shelansky R, Gottscho A, Wagner F, Rouault M, Beliakoff G, et al. High resolution mapping of the breast cancer tumor microenvironment using integrated single cell, spatial and in situ analysis of FFPE tissue. BioRxiv. 2022; p. 2022–10.

5. Mah CK, Ahmed N, Lopez N, Lam D, Monell A, Kern C, et al. Bento: A toolkit for subcellular analysis of spatial transcriptomics data. BioRxiv. 2022; p. 2022–06.

6. Müllner D. Modern hierarchical, agglomerative clustering algorithms. arXiv preprint arXiv:11092378. 2011;.

7. Virtanen P, Gommers R, Oliphant TE, Haberland M, Reddy T, Cournapeau D, et al. SciPy 1.0: fundamental algorithms for scientific computing in Python. Nature methods. 2020;17(3):261–272.

8. SciPy v1.11.4 Manual;. Available from: https://docs.scipy.org/doc/scipy/reference/generated/scipy.cluster.hierarchy.fcluster.html#scipy.cluster.hierarchy.fcluster.
